# Anatomical development of the cerebellothalamic tract in embryonic mice

**DOI:** 10.1101/731968

**Authors:** Daniël B. Dumas, Simona V. Gornati, Youri Adolfs, Tomomi Shimogori, R. Jeroen Pasterkamp, Freek E. Hoebeek

## Abstract

Cerebellar projections to the thalamus are a pivotal connection in cerebello-cerebral interactions. Apart from its role in coordinating sensorimotor integration in the adult brain, the cerebello-thalamic projection has also been implemented in developmental disorders, such as autism spectrum disorders. Although the development of the cerebellum, thalamus and cerebral cortex have been studied in many species, a detailed description of the ontogeny of the mammalian cerebello-thalamic tract (CbT) is currently missing. Here we investigated the development of the CbT at embryonic stages using transgenic *Ntsr1-Cre/Ai14* mice and *in utero* electroporation (IUE) of wild type mice. Wide-field, confocal and 3D light-sheet imaging of immunohistochemical stainings showed that CbT fibers arrive in the prethalamus between E14.5 and E15.5, but only invade the thalamus after E16.5. We quantified the spread of CbT fibers throughout the various thalamic nuclei and found that at E17.5 and E18.5 the ventrolateral, ventromedial and parafascicular nuclei, but also the mediodorsal and posterior complex become increasingly innervated. Several CbT fiber varicosities colocalize with vGluT2, indicating that already from E18.5 the CbT synapse in various thalamic nuclei. Our results contribute to the construction of a frame of reference on the anatomical development of the CbT, which will help to guide future experiments investigating neurodevelopmental disorders.

**Significance statement:** Using various microscopic approaches, we investigated the anatomical development of the fiber tract between the cerebellum and thalamus, one of the major mammalian brain connections. Our results show that in mice cerebellar axons wait outside of the thalamus from embryonic day (E)15.5 until E17.5 before invading the thalamic complex. Cerebellar axons establish vGluT2-positive synapses at E18.5 throughout various thalamic nuclei, each of which subsequently develops its connections with dedicated cerebral cortical regions. Our data thereby advocate the cerebellar influence on the maturation of the thalamus and connected cerebral cortex. This knowledge can help to guide future experiments into neurodevelopmental disorders affecting cerebello-thalamo-cortical networks.

## Introduction

Cerebello-cerebral connectivity is known to be involved in motor activity, as well as several non-motor functions are controlled by long-range cerebellar and cerebral projection neurons (Schmahmann et al., 2009; Manto et al., 2012; Stoodley et al., 2012; Brooks et al., 2015; De Zeeuw and Ten Brinke, 2015). The most direct route from the cerebellum to the cerebral cortex runs through the thalamic complex and it is this cerebello-thalamic (CbT) tract that has been implicated in a wide range of neurological conditions, like rapid onset dystonia, epilepsy and autism spectrum disorder (ASD) (Chen et al., 2014; Kros et al., 2015; Stoodley et al., 2017). Several of these pathologies have a developmental aspect, and thus put a focus on the ontogeny and maturation of the CbT tract. Imaging data from children with cerebellar lesions, which are at high-risk of developing ASD (Wang et al., 2014; Hortensius et al., 2018), revealed concomitant cerebral impairments that have been suggested to be mediated by impairments of CbT connectivity. Yet, the time of onset of the CbT abnormalities in ASD and other neurodevelopmental disorders remains unknown. Despite recent technical advances that allow clinicians to investigate the developing cerebello-cerebral connectivity in very pre-term born children (Pieterman et al., 2016), it remains to be elucidated from which embryonic or fetal stage cerebellar axons reach the thalamic primordium.

In animal models the development of CbT connectivity is equally understudied. Although it is well understood that cerebellar nuclei (CN) neurons are the sole source of the CbT tract, there are few studies available on the development of their axonal connections to the thalamic complex. Some sparse reports on a developing CbT tract in the post-natal opossum (Martin et al., 1987), rat (Cholley et al., 1989; Shirasaki et al., 1995) and a single report in mouse embryo (Hara et al., 2016) describe the timing of CN innervation of midbrain nuclei, such as the parabrachial and red nuclei, and subsequently commence to the thalamic primordium, but no data on the ontogeny of cerebellar innervation of the thalamic complex is currently available. Given that early synaptic afferents have recently been shown to modulate thalamic activity patterns, gene expression profiles and the thalamo-cortical connectivity (Mire et al., 2012; Chou et al., 2013; Moreno-Juan et al., 2017) it is of utmost importance to elucidate at what embryonic stage cerebellar axons start to innervate the developing thalamus. The cerebellum is especially important in this regard, given that in the adult rodent the CbT tract diverges to many first-order and higher-order nuclei, including ventrolateral (VL), ventromedial (VM), centrolateral (CL), posteriomedial (POm), parafascicular (Pf) and mediodorsal (MD) (Bentivoglio and Kuypers, 1982; Shinoda et al., 1985; Sawyer et al., 1994; Teune et al., 2000; Gao et al., 2018; Gornati et al., 2018) each of which has critical periods for growth and maturation of its afferents and efferents. For instance, the somatosensory ventrobasal nuclei receive brainstem afferents from E17.5 (Kivrak and Erzurumlu, 2013) and during the following three weeks thalamic axons initiate connections with various neuronal populations in the developing cerebral cortex (López-Bendito and Molnár, 2003), some of which are transient (Marques-Smith et al., 2016). Disruptions in thalamic and cortical activity and connectivity result in abnormal efferent connections and functional aberrations (e.g., (Tolner et al., 2012; Moreno-Juan et al., 2017). Thus, in the realm of thalamo-cortical connectivity developing activity-dependent and in a time-sensitive fashion, we set out to investigate the ontogeny of the cerebello-thalamic connectivity.

Here we investigated the embryonic development of the CbT tract using transgenic mice in which CbT fibers are tagged with red fluorescent protein (RFP). After confirmation of the presence of the CbT tract in wild type mice by in utero electroporation of CN neurons with a fluorescent tag, we used our transgenic mice to investigate the distribution of CN axons throughout the thalamic primordium between E15.5 and E18.5. Our results reveal that until E16.5 CN axons appear to reside in the prethalamus and then start to innervate thalamic nuclei from its ventro-caudal border. By E18.5 even the rostrally located VL nucleus is innervated by CN axons that show co-labelling with a marker of glutamatergic synapses.

## Materials & Methods

All experiments were performed in accordance with the European Communities Council Directive. Protocols were reviewed and approved by the Dutch national experimental animal committees (DEC) and every precaution was taken to minimize stress and the number of animals used in each series of experiments.

### Mice

To visualize the fibers from the cerebellar nuclei (CN), mice carrying an *Ntsr1-Cre* allele (Gong et al., 2007) (Mutant Mouse Regional Resource Center; Stock *Tg(Ntsr1-cre)GN220Gsat/Mmucd*) were crossed with an *Ai14* reporter line (Madisen et al., 2010) (Jackson laboratories strain 007908) to generate mice expressing RFP in Ntsr1^+^ cells. The lines were genotyped by PCR using dedicated primer sets (5’-GACGGCACGCCCCCCTTA-3’ for *Ntsr1* and 3’-AAGGGAGCTGCAGTGGAGTA-5’ & 5’-CCGAAAATCTGTGGGAAGTC-3’ for wildtype *Ai14* and 3’-CTGTTCCTGTACGGCATGG-5’ & 5’-GGCATTAAAGCAGCGTATCC-3’ for mutant *Ai14*). Only double heterozygous mutants were used for further studies. To investigate the prenatal development of the CbT fibers, we used *Ntsr1-Cre/Ai14* embryos aged from embryonic day (E) 14.5 to E18.5. The morning of the day of vaginal plug detection was counted as E0.5. Dams were deeply anesthetized with isoflurane before extracting the embryos. In total, we used four E14.5, two E15.5, five E16.5, seven E17.5 and six E18.5 embryos. For each age, the embryos were collected from at least two different mothers. There were no gross morphological abnormalities present in any of these embryos. We also used three adult mice (P48-75) for characterization of the *Ntsr1Cre*-expressing CN neurons.

### Tissue preparation for immunohistochemistry

Embryos were collected at E14.5, E15.5, E16.5, E17.5 and E18.5. Those of E14.5 and E15.5 were immediately immersion fixed in 4% paraformaldehyde (PFA); E16.5 and E17.5 were first decapitated before immersion fixation in 4% PFA; and E18.5 brains were immediately dissected in phosphate buffer saline (PBS) over ice before immersion fixation in 4% PFA. All embryo tissue were fixed for 36 hours at 4°C. After fixation, the embryos were cryoprotected in 20% sucrose for at least 3 days at 4°C. After cryoprotection, the embryo tissue was embedded in 22% bovine serum albumin in 7% gelatin solution. The embedded brains were stored at −80°C until sectioning. Sagittal and coronal sections (20 µm) were produced and glass-mounted on chrome alum-gelatin coated slides using a Microm HM560 cryostat (Walldorf, Germany) and then stored at −20°C until processing for immunohistochemistry.

After anesthetizing the adult mice with 0.15 mL pentobarbital (intraperitoneal injection), they were perfused with 4% PFA.The brains of the adult mice were removed after perfusion and post-fixed on a shaker for 2 hrs in 4% PFA at room temperature. After perfusion, the dura mater was removed. Thereafter, the brains were embedded in 12% gelatin/10% sucrose. The gelatin blocks containing the brains were then incubated in 30% sucrose/0.1 M phosphate buffer (PB) overnight at 4°C. Thereafter, the brains were cut into 40 µm thick sections using a Leica SM 2000 R sliding microtome (Nussloch, Germany).

### Immunohistochemistry

For 3, 3-diaminobenzidine (DAB) staining, slides with embryonic sections were immersed in 0.3% H_2_O_2_ in methanol to block endogenous peroxidase activity. After three times of 10 min rinsing with PBS, the cell membranes were permeabilized by immersion in 0.3% Triton in PBS for 60 min. After another three times of 10 min rinsing with PBS, the slides were immersed in a 5% normal horse serum (NHS) in PBS blocking solution for 60 min. Afterwards, the slides were rinsed three times 10 min with PBS and subsequently incubated in primary antibodies in 2% NHS/PBS overnight at 4°C. After three times of 10 min rinsing with PBS, the slides were incubated in secondary antibodies in 2% NHS/PBS for 2 hrs at room temperature. After another three times of 10 min rinsing with PBS, the slides were incubated in avidin biotin complex (ABC) solution for 2 hrs. The ABC solution was prepared 40 minutes before incubation. The ABC solution consists of 0.7% avidin, 0.7% biotin and 0.5% Triton in PBS. After incubation in ABC solution, the slides were rinsed three times 10 min with PBS and two times 10 min with 0.05 M PB. After rinsing, slides were incubated for 15 min in DAB solution, which consisted of 0.665% DAB in 0.1 M PB. The DAB reaction was catalyzed by adding H_2_O_2_ to the solution (final concentration 0.01%) right before immersion. Afterwards, the slides were rinsed three times 10 min in 0.05 M PB. After an additional short rinse in MilliQ, the slides were incubated in thionin for 5 min. Thereafter, the slides were incubated two times 10 min in 96% ethanol, followed by three incubation steps of 2 min in 100% ethanol. Afterwards, the slides were incubated three times 2 min in xyleen and subsequently covered with Permount (Fisher Chemical™ SP15-500) and coverslipped.

To stain embryonic sections with immunofluorescence the slides were first rinsed three times 10 min with PBS and subsequently permeabilized by immersion in 0.3% Triton in PBS for 60 min. The slides were then incubated in 1% sodium dodecyl sulfate in PBS for 5 min to facilitate antigen retrieval. This was followed by times 5 min rinsing with PBS. Thereafter, the slides were immersed in a 5% NHS/PBS blocking solution for 60 min. Afterwards, the slides were rinsed three times 10 min with PBS and subsequently incubated in primary antibodies in 2% NHS/PBS overnight at 4°C. After three times of 10 min rinsing with PBS, the slides were incubated in secondary antibodies in 2% NHS/PBS for 2 hrs at room temperature. After incubation, the slides were rinsed two times 10 min with PBS and 10 min with 0.05 M PB. The slides were then incubated in 1:10000 4’,6-diamidino-2-fenylindool (DAPI) solution for the visualization of cell nuclei. Afterwards, the slides were rinsed two times 10 min with 0.05 M PB and subsequently covered with Mowiol (Sigma Aldrich 4-88) and coverslipped.

For immunofluorescence on adult tissue, the adult brain sections were first rinsed four times 10 min with PBS and subsequently preincubated in 10% NHS / 0.5% Triton / PBS for 60 min. This was followed incubation in primary antibodies in 2% NHS / 0.4% Triton / PBS for 48 hrs at 4°C. Afterwards, the sections were rinsed four times 10 min with PBS and subsequently incubated in secondary antibodies in 2% NHS/ 0.4% Triton/ PBS for 90 min at room temperature. The sections were then rinsed two times 10 min with PBS and 5 min with 0.1 M PB. After rinsing, the sections were incubated in 1:10000 DAPI solution for 10 min. The sections were then rinsed two times for 5 min with 0.1 M PB, after which they were immersed in 5% gelatin / 1% chrome alum / MilliQ. Afterwards, the sections were glass-mounted and coverslipped using Mowiol. Except for the incubation in primary antibody solution, all the steps were performed at room temperature.

### 3DISCO

For this procedure, embryos were collected at E15.5, E16.5, E17.5 and E18.5. The brains of E16.5-18.5 embryos were immediately dissected in phosphate buffer saline (PBS) over ice before immersion fixation in 4% PFA. Younger animals were decapitated and their heads were immediately immersion fixated in 4% PFA. Brains were fixed for 24 hrs at 4°C and then incubated in 0.2% gelatin/ 0.5% Triton/ PBS (PBSGT) for 24 hrs on a shaker (∼70 rounds per min (rpm)) at room temperature. The PBSGT was filtered with a 0.2 µm filter before use. After incubation in PBSGT, brains were incubated in a 0.2 µm filtered primary antibody solution in 0.1% saponin (Sigma Aldrich S-7900)/PBSGT at 37°C on a shaker (∼100 rpm) for 1 week. Afterwards, brains were rinsed six times for 60 min with PBSGT and subsequently incubated overnight in a 0.2 µm filtered secondary antibody solution in 0.1% saponin / PBSGT at 37°C on a shaker (∼100 rpm). Afterwards, brains were rinsed six times for 60 min with PBSGT. Then brains were incubated in 50% tetrahydrofuran (THF; including 250 ppm butylated hydroxytoluene as a stabilizer) (Sigma Aldrich 186562-1L) in H_2_O overnight to start dehydration. Thereafter, brains were incubated for 60 min in 80% THF / H_2_O, then for two times for 60 min in 100% THF. Brains were then incubated in dichloromethane (Sigma Aldrich 270997-1L) for 20 min for clearing the brains, after which they were incubated and stored in dibenzylether (Sigma Aldrich 108014-1KG) at room temperature. From the first THF incubation step onwards, care was taken to have the least amount of air possible in the vials containing the brains (see also (Belle et al., 2014).

### In Utero Electroporation

IUE was performed as described (Matsui et al., 2011). Briefly, timed-pregnant ICR mice with E10.5 embryos were anesthetized with pharmaceutical grade sodium pentobarbital (50 µg per gram body weight). After incision at the abdominal midline, the uterine horns were carefully placed onto a 37 °C pre-warmed phosphate-buffered saline (PBS)-moistened cotton gauze. A flexible fiber optic cable was placed under the uterine horn for visualization of the embryo. After positioning the embryo, a glass capillary was carefully inserted into the fourth ventricle. Embryos were injected with 1 µL of pCAG-EYFP plasmid DNA solution (prepared in TE; 10 mM Tris Base, 1mM EDTA Solution, pH 8.0), 0.1% Fast Green). Embryos were electroporated by inserting a custom-made fine tungsten negative electrode into the fourth ventricle and a custom-made platinum positive electrode into the uterus, placing one hemisphere of the cerebellum between the two electrodes (see (Matsui et al., 2011), for preparation of electrodes). Three square-wave pulses (7V, 100 ms duration) were then delivered with a pulse generator (A-M Systems Model 2100). The uterine horns were returned into the abdominal cavity, the wall and skin were sutured, and the embryos were allowed to continue their normal development until they were sacrificed at E18.5.

### Antibodies

An overview of the antibodies used is presented in **Tables 1** and **2**. For DAB staining, we used a primary rabbit anti-RFP antibody (1:1000, *Rockland*) and a secondary donkey anti-rabbit antibody (1:200, *Jackson*). For immunofluorescence, we used chicken anti-Calbindin (1:500, *Synaptic Systems*), mouse anti-CalbindinD-28 (1:500, *Sigma*), chick anti-GFP (1:200, *Abcam*), goat anti-FoxP2 (1:500, *Santa Cruz*), guinea pig anti-vGluT2 (1:500, *MilliPore*), rabbit anti-RFP (1:1000, *Rockland*), and mouse anti-NeuN (1:1000, *MilliPore*) as primary antibodies. Cy5 anti-chicken (1:200, *Jackson*), Cy3 anti-mouse, FITC anti-chick, (1:200, *MilliPore*), Alexa488 anti-goat (1:200, *Jackson*), Cy5 anti-guinea pig (1:200, *Jackson*), Cy3 anti-rabbit (1:400, *Jackson*), and Alexa488 anti-Mouse (1:200, *Jackson*) antibodies were used as secondary antibodies.

**Table 1:** Primary antibodies

| Host species | Antigen | Concentration | Manufacturer |
| --- | --- | --- | --- |
| Rabbit | Red fluorescent protein | 1:1000 | Rockland 600-401-379 |
| Chick | Green fluorescent protein | 1:200 | Abcam ab13970 |
| Guinea Pig | Vesicular glutamate transporter 2 | 1:500 | MilliP AB2251 |
| Goat | FoxP2 | 1:500 | SC-21069 |
| Chicken | Calbindin | 1:500 | Syn Sys 214006 |
| Mouse | Calbindin D-28 | 1:100 | Sigma CB955 |
| Mouse | NeuN | 1:1000 | MilliP MAB377 |

**Table 2:** Secondary antibodies

| Host species | Target species | Conjugate | Concentration | Manufacturer |
| --- | --- | --- | --- | --- |
| Donkey | Rabbit | Cy3 | 1:400 | Jackson 711-165-152 |
| Donkey | Chick | FITC | 1:200 | Millipore AP194F |
| Goat | Rabbit | Biotin | 1:1000 | Jackson 111-065-144 |
| Donkey | Guinea Pig | Cy5 | 1:200 | Jackson 706-175-148 |
| Donkey | Goat | Alexa488 | 1:200 | Jackson 705-545-147 |
| Donkey | Chicken | Cy5 | 1:200 | Jackson 703-175-155 |
| Goat | Mouse | Cy3 | 1:1000 | Millipore AP124C |
| Donkey | Mouse | Alexa488 | 1:200 | Jackson 715-545-150 |

For 3DISCO, we used chicken anti-Calbindin (1:500, *Synaptic Systems*), goat anti-FoxP2 (1:500, *Santa Cruz*) and rabbit anti-RFP (1:000, *Rockland*) as primary antibodies. Cy5 anti-chicken (1:200, *Jackson*), Alexa488 anti-goat (1:200, *Jackson*) and Cy3 anti-rabbit (1:400, *Jackson*) antibodies were used as secondary antibodies.

### Imaging

An overview of the microscopes used is presented in **Table 3**. Light microscopy pictures of DAB stained slides were made using a Nanozoomer 2.0-HT (Hamamatsu, Japan) at 40X magnification captured using a 20X objective with an NA of 0.75. The pixel size was 230 nm × 460 nm. Confocal microscopy pictures were taken with a Zeiss LSM700 Meta (Carl Zeiss Microscopy, LLC, USA), an Olympus FV3000 (Olympus, Tokio, Japan) and an Opera Phenix^TM^ HCS system (Perkin Elmer, Hamburg, Germany). Confocal pictures on the Zeiss LSM700 Meta were taken with a 63X oil Plan-Apochromat lens with an NA of 1.4. Z-stacks were taken with a voxel size of 50 * 50 * 150 nm, a pinhole of 1 Airy unit and bit depth of 8-bits. Signal-to-noise ratio was improved by 4X line averaging. For the different fluorophores, the following lasers were used: 405 nm for DAPI, 488 nm for Alexa488, 543 nm for Cy3 and 633 nm for Cy5. The confocal pictures IUE tissue were acquired on Keyence (BZ-X800, Itasca, IL, USA) and taken with a 10X lens with an NA of 0.45. The confocal pictures on the Opera Phenix^TM^ HCS system were taken with a 20X lens with an NA of 0.4. The pixel size was 598 nm × 598 nm. The bit depth was 16-bits. For the different fluorophores, the following lasers were used: 488 nm for Alexa488, 561 nm for Cy3 and 640 nm for Cy5. Z-stacks consisting of two 10 µm spaced slices were taken to correct for shifts in the z-axis. Before further analysis, a maximum intensity projection of the images was produced, which was subsequently converted to 8-bits.

**Table 3:** Laser and filters per microscope per dye

|  | Light-sheet<br>Ultramicroscope II |  | Opera Phenix™ HCS |  | Zeiss LSM700 Meta |  |  |
| --- | --- | --- | --- | --- | --- | --- | --- |
| Dyes | Laser | Filters | Lasers | Filters | Lasers | Filters | Beamsplitter |
| Alexa 488 | - | - | 488nm | 500/50 | 488nm | 0/590 | 520 |
| Cy3 | 561nm | 615/40 | 561nm | 570/30 | 555nm | /640 | 600 |
| Cy5 | - | - | 640nm | 650/60 | 633nm | 640/ | 630 |

The 3DISCO cleared brains were imaged using LaVision biotec light sheet microscope Ultramicroscope II (LaVision biotec, Bielefeld, Germany). The microscope consists of an Olympus MVX-10 Zoom Body (0.63-6.3x) equipped with an Olympus MVPLAPO 2X objective lens, which includes dipping cap with 6 mm working distance. Images where taken with a Neo sCMOS camera (Andor) (2560 × 2160 pixels; Pixel size: 6.5 × 6.5 µm). Samples were scanned with two-sided illumination, a sheet NA of 0.148348, which results in a 5 µ m thick sheet. We applied a step-size of 2.5 µm using the horizontal focusing light-sheet scanning method and contrast blending algorithm. The effective magnification for overview pictures was 3.2X and for detailed images 12.6X. The lens had a NA of 0.5 and bit depth was 16-bits. Overview pictures had a voxel size of 2030 nm × 2030 nm × 2500 nm and detailed pictures had a voxel size of 515.9 nm × 515.9 nm × 2500 nm. The following laser filter combination was used to image Cy3: Coherent OBIS 561-100 LS Laser with 615/40 filter.

### Delineation and intensity measurements

Any images containing structural artefacts (e.g. freezing artifacts and cutting artifacts) were discarded. The delineation of anatomical regions and measurements were performed with FIJI software (version 1.51). To delineate the thalamic nuclei across the different ages studied, we used the chemoarchitectonic atlas of the developing mouse brain (Jacobowitz and Abbott, 1998), the atlas of the prenatal mouse brain (Schambra et al., 1992), the atlas of the developing mouse brain (Paxinos, 2007), the Allen brain atlas (Thompson et al., 2014), and descriptive studies of FoxP2 expression in developing mice (Ferland et al., 2003; Vargha-Khadem et al., 2005; Thompson et al., 2014; Nagalski et al., 2016). With FoxP2, we could delineate Pf, MD, VB, POm, VM and midline (ML) nuclei, the latter of which we defined as centromedial, paraventricular, intermediodorsal, reunions, retroreuniens, intermediodorsal, rhomboid, xyphoid, retroxyphoid, interanteromedial, anteromedial and posteromedian nuclei of the thalamus. When combining FoxP2 with calbindin staining, we could delineate the VM and the ML more specifically than with FoxP2 alone (Bodor et al., 2008). In addition to these markers, we also used the atlases described in the materials and methods section. The border between VL and the intralaminar nuclei could not consistently be accurately delineated in all slices. Therefore, the size of the nucleus might be slightly underestimated in some instances. Moreover, at E16.5, the border between the VL and the LP could not be delineated. We therefore did not measure the VL at this age. The delineation of thalamic nuclei was performed in the fluorescent channels representing the FoxP2 and calbindin expression, so that the researcher was blinded for the location of RFP signal, which represents the location of *Ntsr1Cre-Ai14*-positive axons. For further measurements, nuclei were delineated with the ROI manager in FIJI. If a nucleus could not be delineated in two or more sequential sections, the data for this particular nucleus of that mouse was excluded.

To measure the amount of Cy3 signal in each nucleus, the mean plus two times the standard deviation of the histogram was used as threshold to binarize the image into background and foreground. To measure the area of detected objects within each nucleus (A_detected_) we used the ‘analyze particle’ function.

### Colocalization

Colocalization of vGluT2 and RFP signals was determined using FIJI in the 63X z-stacks of the different thalamic nuclei. After applying user defined thresholds in the separate channels of the raw z-stacks, sites of putative localization were subjectively identified by the appearance of structures showing the presence of both colors and then deconvolved using Huygens (Scientific Volume Imaging). After deconvolution, a user defined threshold was applied to confirm the colocalization, which was defined as a complete overlap of vGluT2 and RFP structures (see also (Gornati et al., 2018)). In short, vGluT2 positive terminals’ volume in VL and VM was measured using a custom-written Fiji macro. This macro used manual thresholding and analyze particles to acquire masks of the vGluT2 and RFP channels on which the Image Calculator was used to acquire an image showing only the overlap between vGluT2 and RFP. The volume thereof was then determined with 3D object counter.

### Statistical Analysis

The fluorescence of a whole thalamic nucleus was calculated for each mouse by summing the A_detected_ and the A_delineated_ of all the sections containing the nucleus in question, yielding the ∑A_detected_ and ∑A_delineated_, respectively, per nucleus per hemipshere. Dividing ∑A_detected_ by ∑A_delineated_ gives the sum of the relative area occupied by RFP^+^ fibers, expressed by the percentage of Summed Area Occupied (pSAO). For each mouse, data from both hemispheres were averaged. Thereafter, the average pSAO per nucleus was calculated for E16.5, E17.5 and E18.5. For each nucleus, significant differences between the ages were first tested with a Kruskal-Wallis (K-W) test (degrees of freedom = 2). When a significant difference was found, Dunn’s post-hoc test was used to compare pairwise between the age groups. For the latter test, a Šidák corrected *p*-value of 0.0127 was used as threshold for significance. To compare the data of nuclei gathered from only two ages we used a Mann Whitney U test with a Šidák corrected *p*-value of 0.017 as threshold for significance. To compare the data of MD, POm and VB combined with VM, VL and Pf separately, we used a Mann Whitney U test with a Šidák corrected *p*-value of 0.0127. Šidák corrected p-values are calculated using 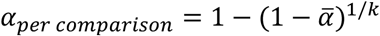, where 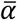 is the overall significance level, which was chosen to be 0.05, and k was chosen as the amount of times a same dataset was analyzed, or the amount of comparisons in the case of a Dunn’s post-hoc test.

The relative amount of RFP^+^ fibers per section was calculated by dividing the A_detected_ by the A_delineated_. These values were then plotted against the relative caudal-to-rostral distance. This distance was calculated as a linear scale from 0 to 100, with 0 indicating the most caudal section in which a particular nucleus was delineated and 100 indicating the most rostral section in which that nucleus was delineated. To determine whether there was caudal-to-rostral gradient of the relative amount of RFP^+^ fibers, a Spearman’s rank correlation coefficient ‘rho’ was calculated per nucleus per age. To measure differences in correlation between groups, the Fisher’s Z-transformation was used, after which a Z-test was conducted for pairwise comparison. Since the same dataset was used for the former analysis, a Šidák corrected p-value of 0.017 was used as threshold for significance.

## Results

### Characterization of the Ntsr1-Cre/Ai14 mouse line

To study the embryonic development of the CbT tract, we crossed the *Ntsr1-Cre* and *Ai14* mouse lines. Offspring heterozygous for both constructs is characterized by RFP^+^ expression in previously characterized large diameter CN neurons (**Fig. 1A-D**) (see also (Houck and Person, 2015)). To assess the distribution of Cre expression in the cerebellum we assessed the proportion of RFP^+^ CN cells and the number of NeuN^+^ cells in the CN (**Fig. 1E**). Since there are non-CN neurons migrating through the subcortical part of the cerebellum at embryonic stages (Englund et al., 2006; Leto et al., 2015), we quantified the RFP^+^ CN population in adult mice (**Table 4**). Most of these RFP^+^ neurons reside in the interposed nucleus and appeared larger than RFP^−^ somata. In contrast to the CN, no RFP^+^-neurons were found in the nearby vestibular nuclei or in the cerebellar cortex (data not shown).

**Figure 1.**
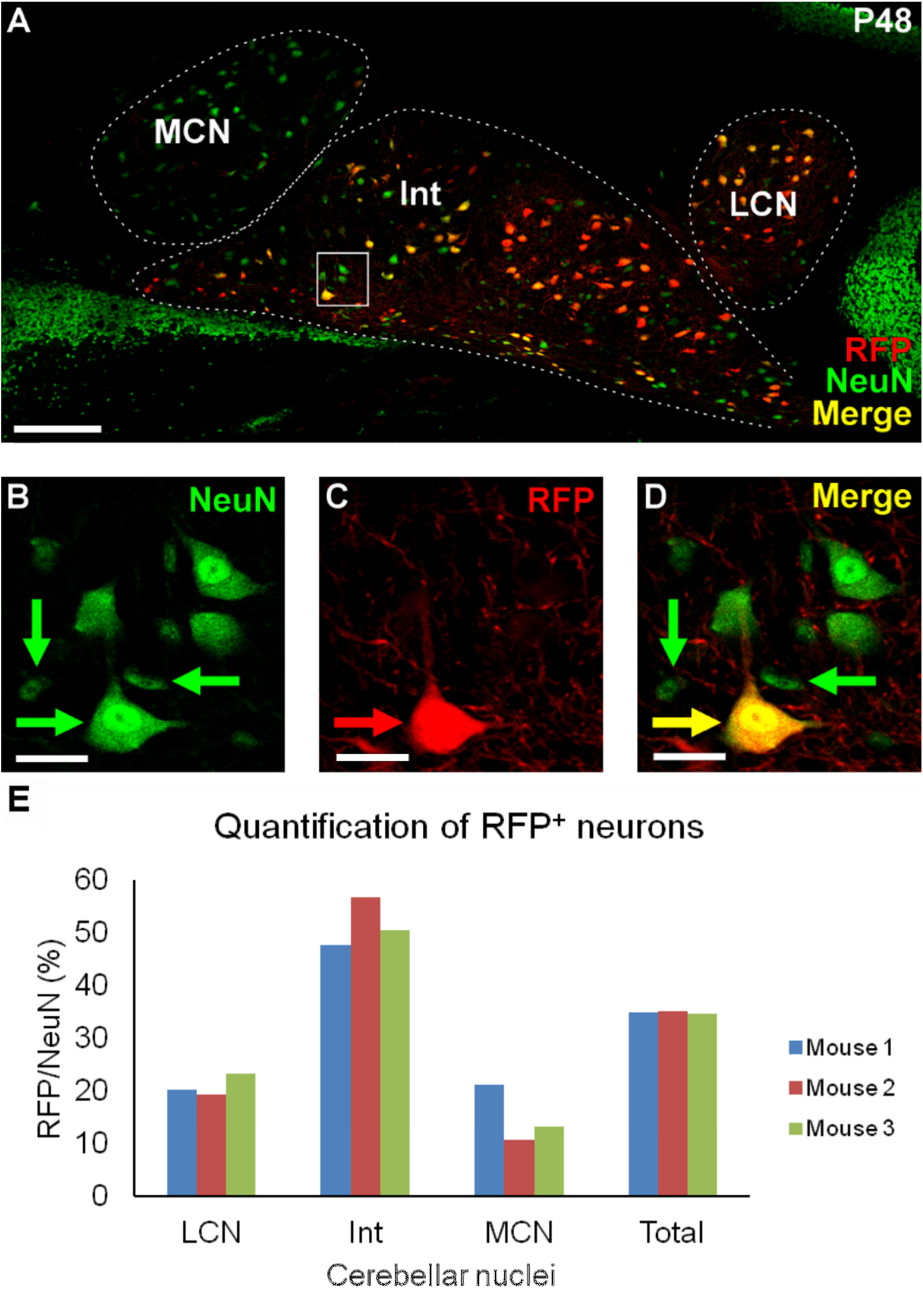
Example of NeuN and RFP staining in adult CN. A) 20x tilescan of a coronal P48 cerebellar slice, zoomed in on the left CN. Stainings: RFP in red, NeuN in green, yellow indicating colocalization of these two stainings. B-D) Zoom in of boxed region in A. B) NeuN stained cells. C) RFP stained cells and fibers. D) Merge of B and C. Note that the RFP^+^ neurons are relatively big and all small neurons are RFP^−^. E) A bar plot representation of the quantification of RFP^+^ cells as a proportion of NeuN^+^ cells (N = 3 mice). Scale bars: A = 150 µ m, B-D = 25 µ m. Int = interposed nucleus; LCN = lateral cerebellar nucleus; MCN = medial cerebellar nucleus; RFP = red fluorescent protein.

**Table 4:** Relative amount of RFP^+^ neurons (RFP / NeuN in %) per cerebellar nucleus and the relative amount of RFP^+^ neurons in each nucleus compared to the total amount of RFP^+^ CN neurons (RFP / totalRFP in %). RFP = red fluorescent protein; int = interposed nucleus; LCN = lateral cerebellar nucleus; MCN = medial cerebellar nucleus.

|  | Mouse 1 |  | Mouse 2 |  | Mouse 3 |  |
| --- | --- | --- | --- | --- | --- | --- |
|  | RFP/NeuN<br>in % | RFP/totalRFP | RFP/NeuN<br>in % | RFP/totalRFP | RFP/NeuN<br>in % | RFP/totalRFP |
| LCN | 33.3 | 22.1 | 22.5 | 13.3 | 23.2 | 16.1 |
| Int | 56.6 | 64.8 | 64.6 | 77.5 | 50.4 | 74.3 |
| MCN | 19.0 | 13.1 | 12.7 | 9.2 | 13.4 | 9.6 |
| Total | 40.0 | 100 | 39.7 | 100 | 34.6 | 100 |

During embryonic development, RFP^+^ neurons are also found outside of the CN (**Fig. 2**). Apart from some sparse RFP^+^ neurons throughout various regions that do not project primarily to the cerebellar-receiving thalamic nuclei, i.e. the posterior and lateral hypothalamus, hippocampus, lateral geniculate nucleus, medial reticular formation, superior colliculus and the Rim, the only other region that showed pronounced RFP^+^ expression at E18.5 was layer VI of the cerebral cortex (Gong et al., 2007; Olsen et al., 2012). Since cortical layer VI neurons project throughout the thalamic complex from E17.5 onwards (Jacobs et al., 2007), we investigated the relative location of their axons compared to RFP^+^ axons from CN neurons. We found that at E18.5 RFP^+^ axons from cortical layer VI via the internal capsule reached the VB nucleus, but remained lateral of VM (**Fig. 3**). The RFP^+^ CN axons reside more medially in the bundle that enters the thalamic complex from the mesencephalic and subthalamic regions. To confirm the presence of cerebellar axons in the thalamic primordium at E18.5, we labelled CbT fibers using IUE transfection with a EYFP-construct at E10.5. Apart from a dense population of transfected cerebellar neurons and sparse labelling in the brainstem (**Fig. 4A,B**), the rest of these E18.5 brains solely contained labelled axons with dense diencephalic projections. Qualitatively, the CbT tract labelled via IUE transfection is similar to the presumptive CbT labelled in our *Ntsr1-Cre/Ai14* embryos at E18.5 (**Fig. 4C**), in that ventrobasal thalamic nuclei are largely avoided by CN axons in contrast to, for instance, VM. These results from IUE transfection of wildtype hindbrain neurons indicate that at late embryonic stages RFP^+^ axons in *Ntsr1Cre-Ai14* thalami predominantly represent CbT fibers.

**Figure 2.**
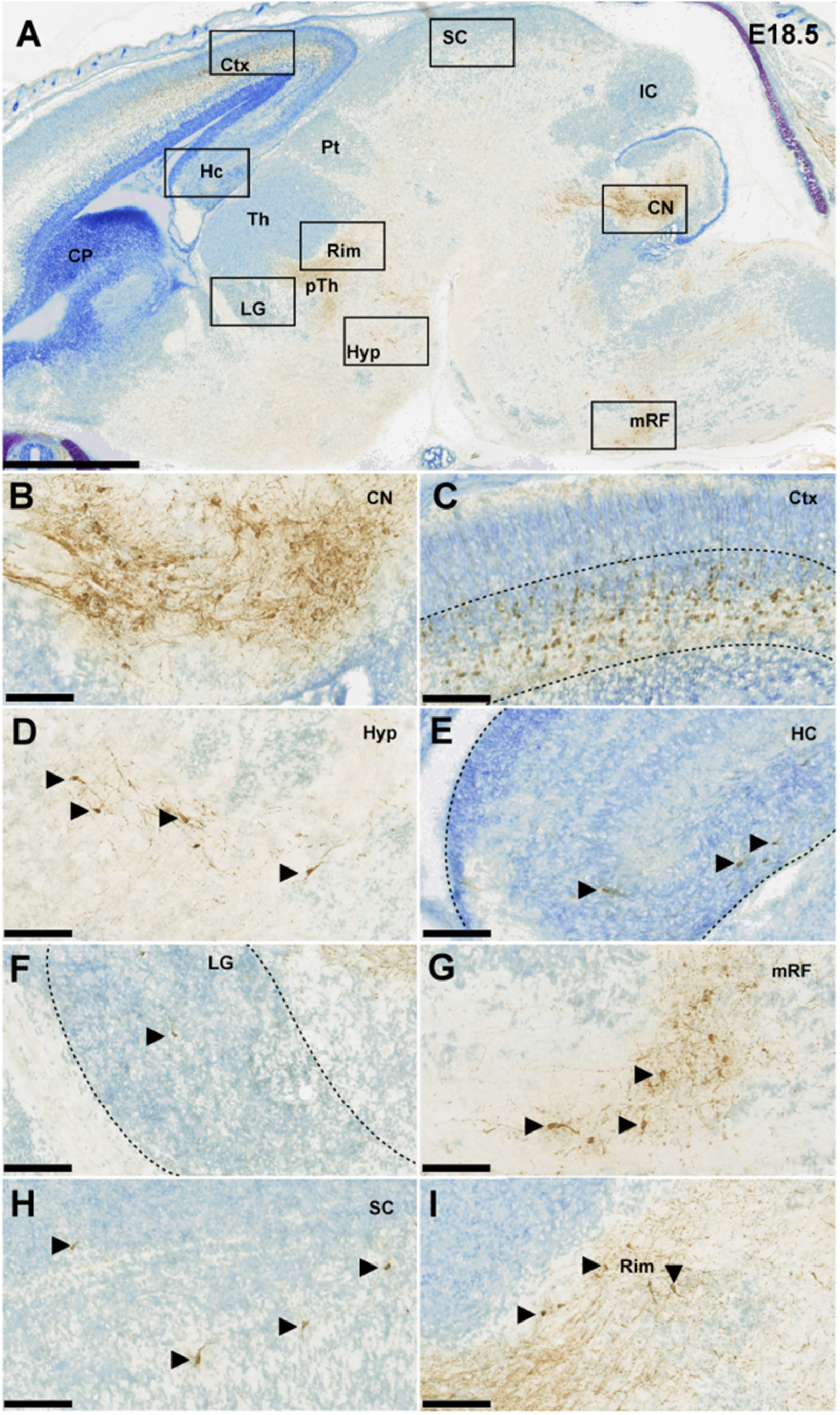
Ntsr1^+^ cells DAB-stained in an E18.5 embryonic mouse brain; counterstained with thionin. A) Sagittal section showing all the regions in which Ntsr1 is expressed. Scale bar = 1000 µm B-H) 20x zoom in pictures of the CN, the Ctx, the Hyp, the Hc, the LG, the mRF and the SC, respectively. I) 20X zoom in image of a region at the interface between Thalamus and pTh with Ntsr1^+^ cells presumably originating from the Rim. Scale bar = 100 µm. CP = caudate putamen, Ctx = cerebral cortex, Hc = hippocampus, Hyp = hypothalamus, IC = inferior colliculus, LG = lateral geniculate nucleus, mRF = medial reticular formation, Pt = pretectum, pTh = prethalamus, SC = superior colliculus, Th = thalamus.

**Figure 3.**
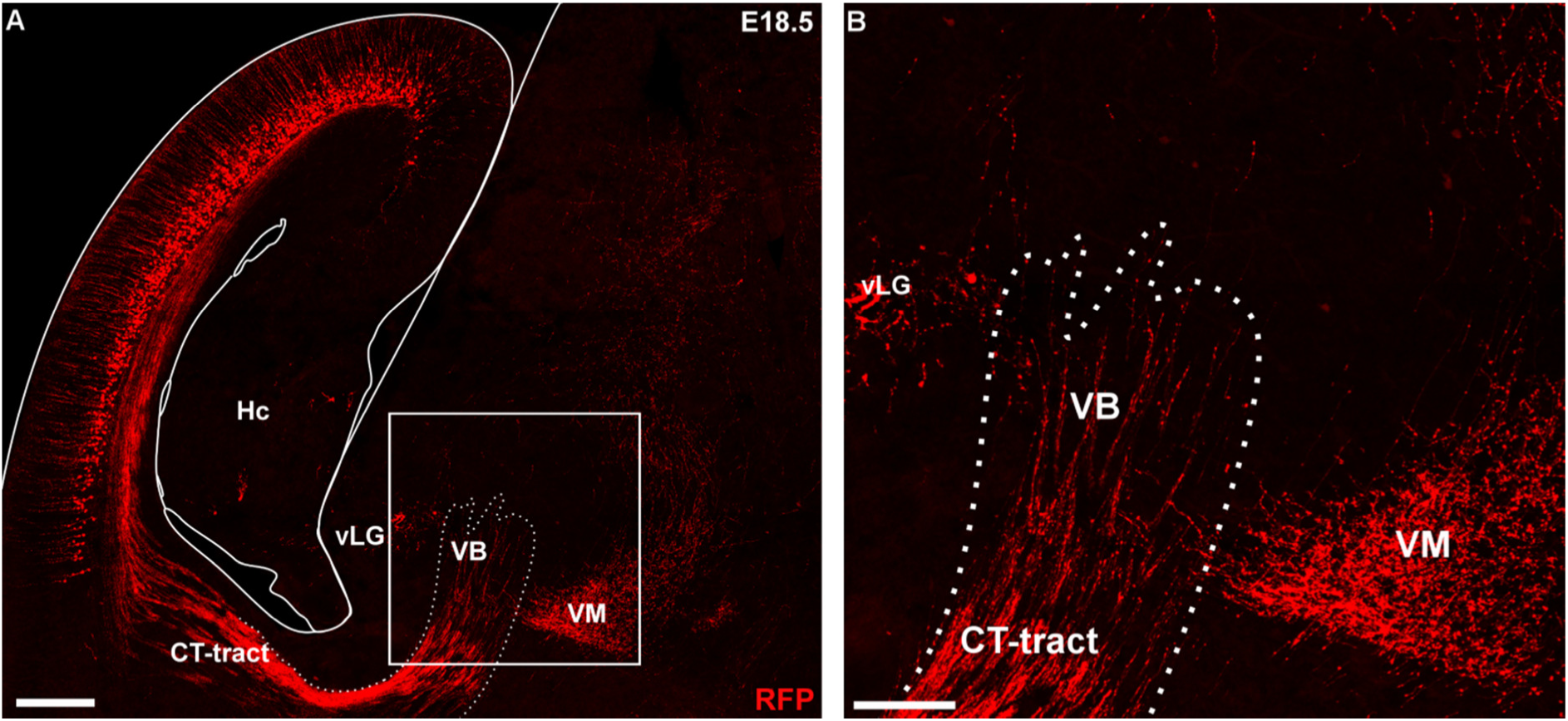
RFP^+^ cortical fibers entering the thalamus, specifically the ventrobasal thalamic nucleus, at E18.5. A) Zoomed out confocal tile scan showing the RFP^+^ CT fibers and another, separate, RFP^+^ fiber bundle, presumably originating from the CN B) Zoom in of inset in A. Scale bars: A = 250 µm, B = 100 µm. CN = cerebellar nuclei; CT = corticotalamic; Hc = hippocampus; RFP = red fluorescent protein; vLG = ventral lateral geniculate nucleus; VB = ventrobasal nucleus; VM = ventromedial nucleus.

**Figure 4.**
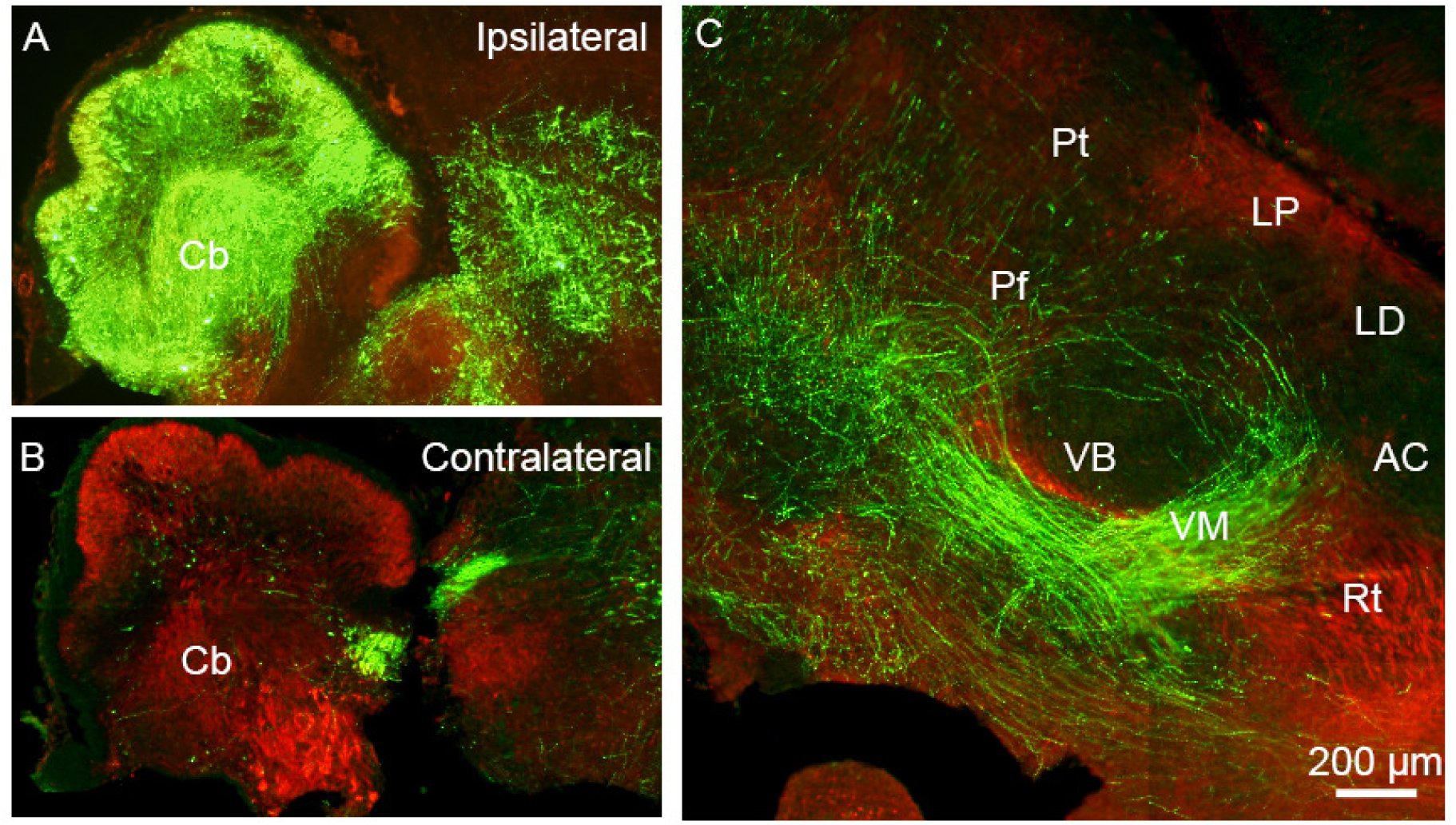
EYFP^+^ fibers of IUE transfected cerebellum are present in the thalamus at E18.5. A, B) EYFP^+^ signal in the transfected ipsilateral cerebellum. C) EYFP^+^ fibers in the thalamic complex show selective innervation of particular nuclei. Note that in this sagittal section several CbT-innervated nuclei are outside the field of view, i.e. on a different level of the medio-lateral axis. AC = anterior complex; Cb = Cerebellum; CbT = cerebellothalamic tract; EYFP = enhanced yellow fluorescent protein; LD = lateral dorsal nucleus; LP = lateral posterior nucleus; Pf = parafascicular nucleus; Pt = pretectum; Rt = reticular nucleus; SC = superior colliculus; VB = ventrobasal nucleus; VM = ventromedial nucleus.

### RFP^+^ axons of the CbT tract reside in prethalamus until E16.5

To determine at which age RFP^+^ CN axons arrive at the thalamic primordium, we used a combination of light-sheet imaging of 3DISCO-treated brains and confocal and light-microscopy of histological sections (**Fig. 5**). At E14.5 the CbT did not yet reach diencephalic structures (data not shown; (Hara et al., 2016). At E15.5 and E16.5 the CbT is positioned dorsally in the mesencephalic curvature and has progressed beyond the red nucleus to reach the prethalamus (**Fig. 5C,D,G,H**). Yet, in E16.5 brains we found no evidence for CbT fibers invading the thalamic complex.

**Figure 5.**
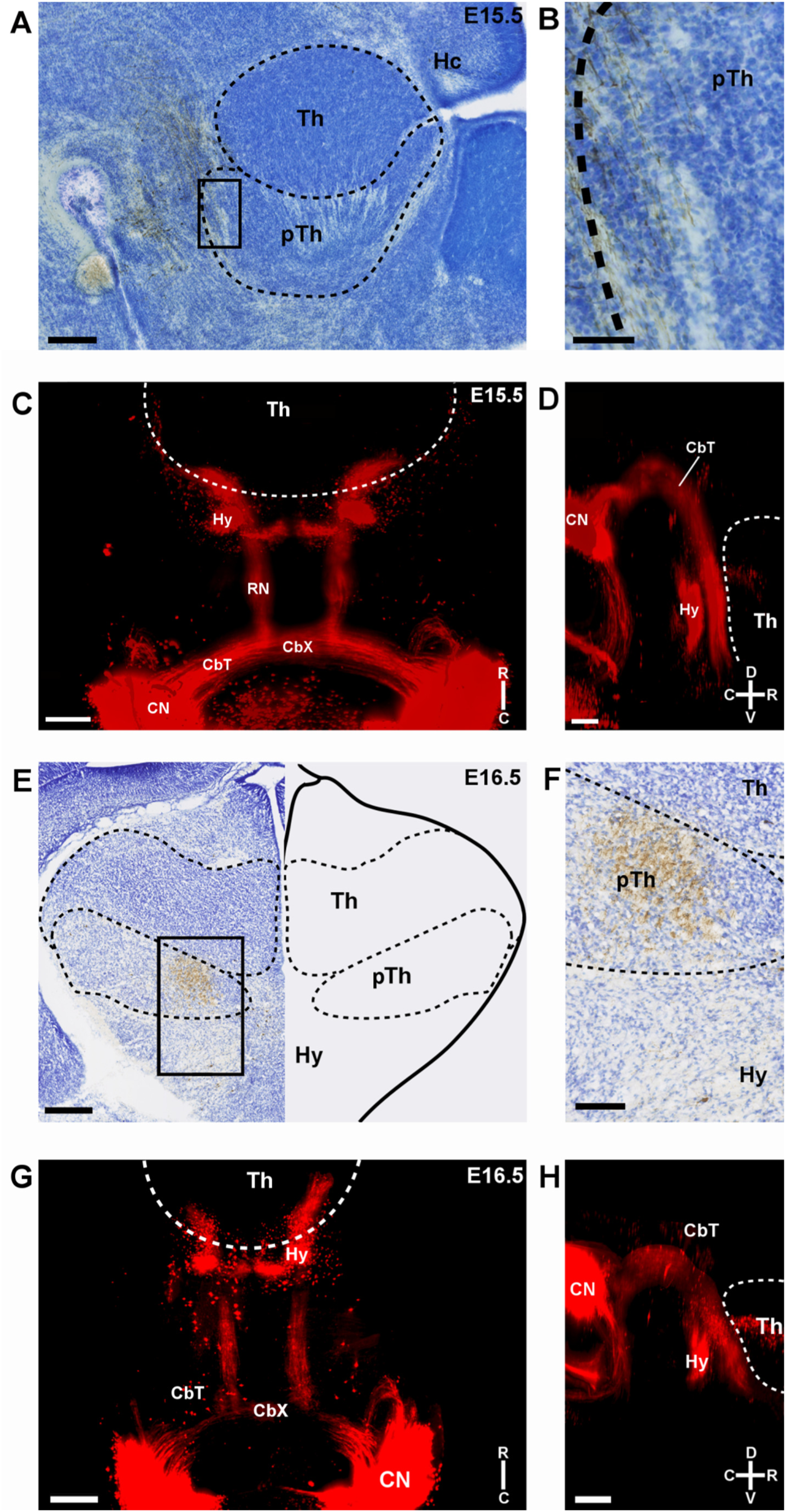
RFP^+^ fibers in the prethalamus at E15.5 and E16.5. A-B) A sagittal section at E15.5 showing the presence of DAB-stained RFP^+^ fibers in the pTh and absence of RFP^+^ fibers in the thalamus at this age. Counterstained with thionin. C-D) Respectively horizontal and sagittal maximum intensity projections of an RFP-stained, 3DISCO cleared brain, imaged with light-sheet microscopy. Note that some RFP-staining appears to reside within thalamus, but actually is located outside of the thalamus – this comes about because of a 2D representation of a 3D image. E-F) Coronal section of an E16.5 mouse brain with inset, respectively, showing presence of DAB-stained RFP^+^ fibers in the pTh and absence of RFP^+^ fibers in the thalamus at this age. Counterstained with thionin. G-H) Respectively horizontal and sagittal maximum intensity projections of an RFP-stained, 3DISCO cleared brain, imaged with light-sheet microscopy. Scale bars: A = 100 µm, B = 50 µm, C and D = 100 µm, E = 250 µm, F = 100 µm, G and H = 250 µm. Compasses in bottom right of each 3DISCO panel indicating directions of rostral (R), dorsal (D), caudal (C) and ventral (V) sides. CbT = cerebellothalamic tract; CbX = decussation of the CbT; CN = cerebellar nuclei; Hy = hypothalamus; pTh = prethalamus; RFP = red fluorescent protein; RN = red nucleus; Th = thalamus.

### CbT RFP^+^ axons progressively innervate specific nuclei in thalamus from E17.5

From E17.5 the CbT commences to innervate the thalamic nuclei (**Fig. 6A-E**). To establish which nuclei are invaded by RFP^+^ axons we combined immunofluorescent staining for FoxP2 and calbindin-d28K on alternating slices, combined with RFP-staining to identify the neuronal populations that form the separate nuclei (**Fig. 6F**). We found from E17.5 that the bundle of RFP^+^ axons extended from the prethalamus into the nearby VM nucleus and diverged towards other nuclei (**Fig. 6G,H**).

**Figure 6.**
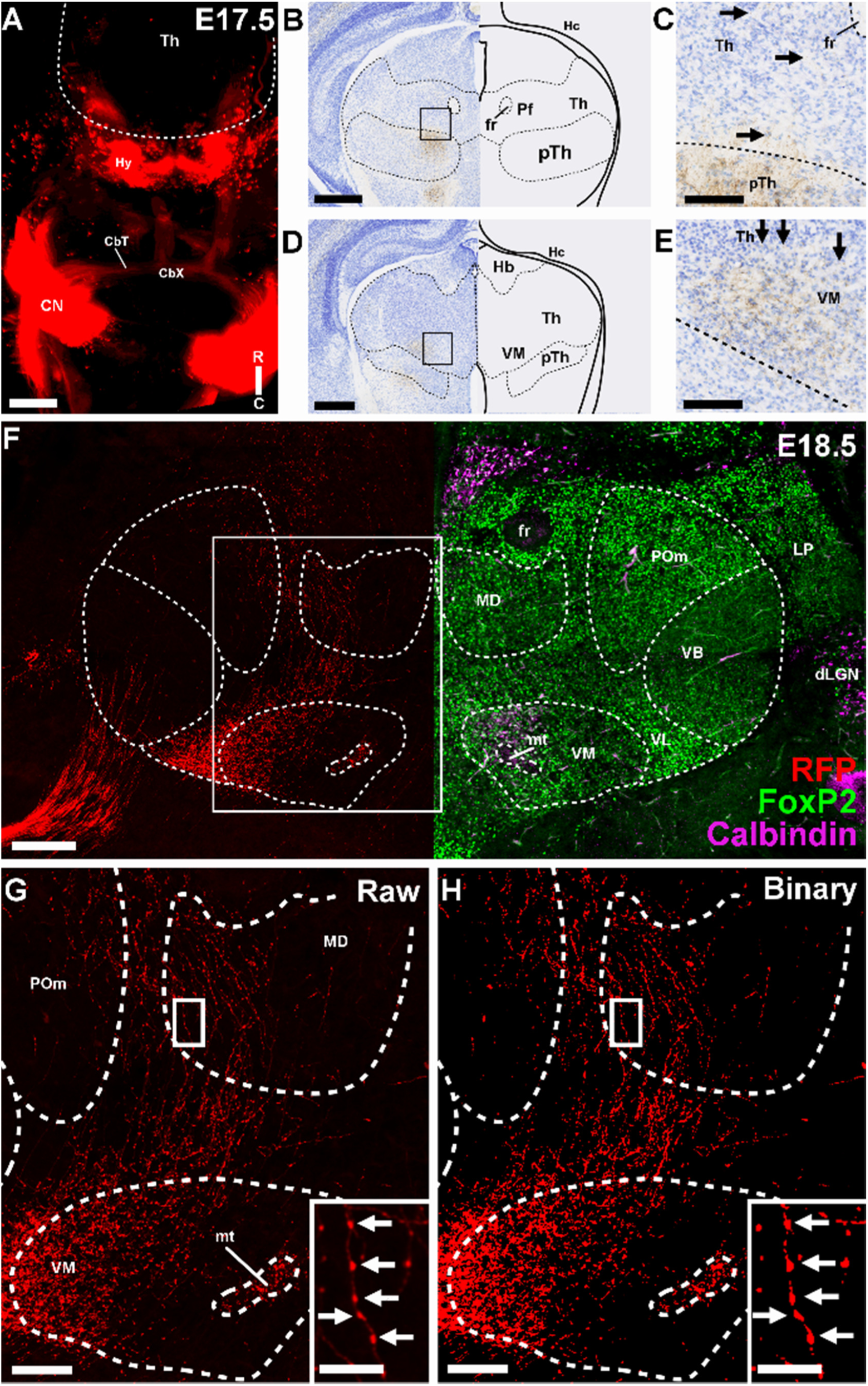
RFP^+^ fibers in the diencephalon at E17.5 and E18.5. A) Horizontal view of a maximum intensity projection of a 3DISCO cleared E17.5 mouse brain. No visible change has occurred compared to E16.5 at this level of magnification. Compass in bottom right indicating directions of rostral (R) and caudal (C) sides B-E) Coronal section of an E17.5 mouse brain, with insets. Fibers can be seen in the Th, though they are much less bundled than in the medial pTh. F) Coronal section of an E18.5 mouse brain stained for FoxP2 (in green), Calbindin (Magenta) and RFP (in red) showing an example of delineation of the VM, POm, MD, VB, VL, LP, dLGN, fr, and the mt. G) Zoom in of boxed region in F. In the inset, a single axon with multiple putative boutons is visible. Arrows indicate the putative boutons. H) Binarized version of G after applying the threshold. Inset is a binarized version of the inset in H. Scale bars: A = 100 µm; B and D = 250 µ m; C and E = 100 µ m; F = 200 µm; G = 100 µm, insets = 25 µm. CbT = cerebellothalamic tract; CbX = decussation of the CbT; CN = cerebellar nuclei; dLGN = dorsal lateral geniculate nucleus; fr = fasciculus retroflexus; Hb = Habenula; Hy = hypothalamus; LP = lateral posterior nucleus; MD = mediodorsal nucleus; mt = mamillothalamic tract; Pf = Parafascicular nucleus; POm = posteromdeial nucleus; VB = ventrobasal nucleus; VL = ventrolateral nucleus; VM = ventromedial nucleus.

To assess the innervation of the individual nuclei by RFP^+^ axons, we selected all sections available for the identified nuclei and summed the percentage of the area that was RFP^+^ (see also **table 5**). At E17.5 the surface covered by VM, VL and Pf, i.e., nuclei which receive dense CN innervation in the adult mouse brain (Gornati et al., 2018), was ≥1% RFP^+^ (VM: 4.85 ± 4.13%; VL: 2.20 ± 1.41%; Pf: 1.06 ± 0.41%). In E18.5 tissue the surface of RFP-signal further increased: in VM we found 14.7 ± 2.1% of the section’s surface to be RFP^+^, in VL 9.56 ± 2.95% and in Pf 3.30 ± 1.55% (**Fig. 7A-C**). Also beyond the VM, VL and Pf nuclei we found that during the final days of embryonic development the relative surface of RFP^+^ signal within MD and POm tended to increase (**Fig. 7D-G**). At E18.5 the MD, POm and VB did not differ significantly from one another, but compared to these nuclei the surface was found to be significantly less in ML. In addition, the relative surface of RFP-signal within MD, POm and VB, at E18.5 combined (1.08 ± 0.54%) remained below the RFP fluorescence levels found in VM (p = 0.0036) and VL (p = 0.0036) but not Pf (p = 0.015).

**Figure 7.**
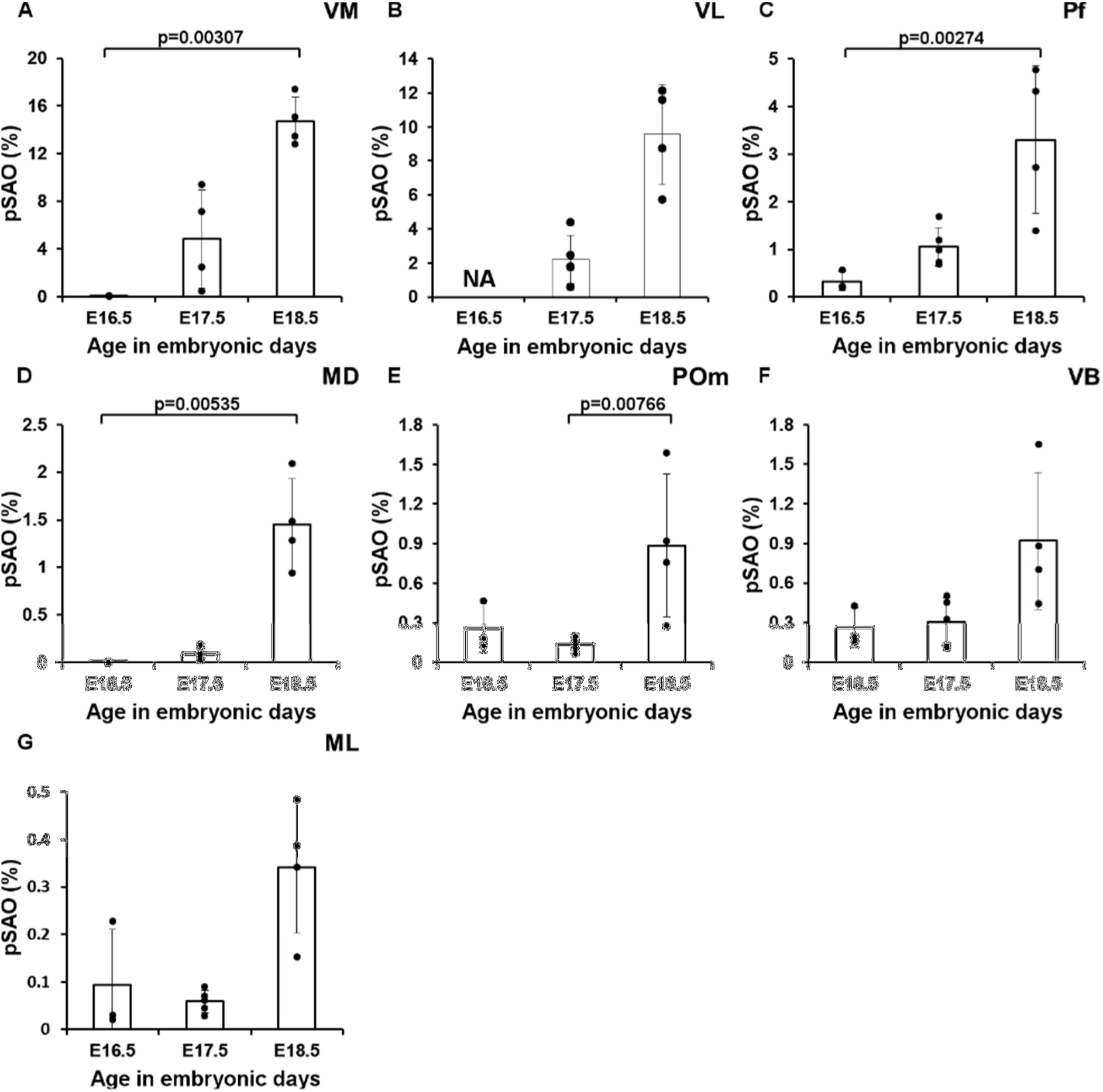
RFP signal quantification in the thalamic nuclei. Percentage of summed area occupied by above-threshold RFP-signal (pSAO) in A-G. A) VM (E16.5, n=3, E17.5, n=4, E18.5, n=4), B) VL (E16.5, n=0, E17.5, n=4, E18.5, n=4) C) Pf (E16.5, n=3, E17.5, n=5, E18.5, n=4), D) MD (E16.5, n=2, E17.5, n=5, E18.5, n=4), E) POm (E16.5, n=3, E17.5, n=5, E18.5, n=4) F) VB (E16.5, n=3, E17.5, n=5, E18.5, n=4), G) ML (E16.5, n=3, E17.5, n=5, E18.5, n=4). Since the VL could not reliably be delineated at E16.5, as the border between VL and the intralaminar nuclei could not consistently be accurately delineated in all slices, we measured the pSAO in this nucleus at E17.5 and E18.5. Columns represent the mean; error bars represent ±SD; dots represent individual data points. MD = mediodorsal nucleus; Pf = Parafascicular nucleus; POm = posteromdeial nucleus; VB = ventrobasal nucleus; VL = ventrolateral nucleus; VM = ventromedial nucleus

**Table 5:** pSAO values (in %) of the investigated thalamic nuclei and the Z- and p-values of the differences between the age groups of each of these nuclei. Šidák corrected *p*-values of 0.0127 were used for all nuclei, except VL for which a Šidák corrected *p*-value of 0.017 was chosen. Significant differences are indicated in bold letters. For VB there was no post-hoc test since the Kruskal Wallis analyses showed no significant difference. MD = mediodorsal nucleus; ML = midline nucleus; Pf = parafascicular nucleus, POm = posteromedial nucleus; pSAO = percentage of summed area occupied; VB = ventrobasal nucleus; VL = ventrolateral nucleus; VM = ventromedial nucleus.

|  | E16.5 | E17.5 | E18.5 | E16.5 vs. E17.5 |  | E16.5 vs. E18.5 |  | E17.5 vs. E18.5 |  |
| --- | --- | --- | --- | --- | --- | --- | --- | --- | --- |
|  | pSAO | pSAO | pSAO | Z | P | Z | p | Z | p |
| VM | 0.0480% | 4.85% | 14.7% | -1.38 | 0.167 | -2.96 | **0.00307** | -1.71 | 0.0881 |
| VL | - | 2.20% | 9.56% | - | - | - | - | - | 0.02 |
| MD | 0.00147% | 0.0590% | 1.45% | -1.26 | 0.207 | -2.79 | **0.00535** | -2.02 | 0.0431 |
| Pf | 0.330% | 1.06% | 3.30% | -1.60 | 0.111 | -3.00 | **0.00274** | -1.67 | 0.0940 |
| ML | 0.0930% | 0.0586% | 0.342% | -1.77 | 0.859 | -2.15 | 0.0317 | -2.25 | 0.0242 |
| VB | 0.263% | 0.305% | 0.919% | - | - | - | - | - | - |
| POm | 0.0261% | 0.137% | 0.885% | 0.836 | 0.403 | -1.54 | 0.123 | -2.67 | **0.00766** |

To describe the development of the CbT in more detail, we focused on the VM, VL and Pf nuclei and analysed the caudal-to-rostral gradient of the relative amount of RFP^+^ fibers (**Fig. 8**). We calculated the Spearman’s correlation value rho (r) for E17.5 and E18.5 tissue pooled from various embryos (see methods) and found that in all these nuclei the fluorescence was relatively higher in the most caudal sections. In VM at E17.5 we found significant negative correlation between the rostro-caudal level and the relative amount of RFP^+^ fibers (r=−0.550, df=48, p=3.57*10^−5^) (**Fig. 8A**). Also in E18.5 embryos we found this negative correlation value for VM (pooled r=−0.790, df=57, p=1.06*10^−13^) (**Fig. 8B**). When comparing the Fisher’s Z-transform of the correlation values for VM, we found that the correlation was significantly stronger at E18.5 than at E17.5 (−0.790 vs. −0.550, Z=−2.29, p=0.0226; for E18.5 vs. E17.5, respectively). In the VL at E17.5, there was a negative correlation between the rostro-caudal level and the relative amount of RFP^+^ fibers (r=−0.711, df=44, p=3.11*10^−8^) (**Fig. 8C**). At E18.5 this correlation seemed weaker (r=−0.591, df=50, p=4.06^−6^) (**Fig. 8D**); however, when comparing the Fisher’s Z-transform of the correlation values this correlation did not differ significantly from that of the E17.5 animals (Z=1.01, p=0.844). In the Pf we found the same negative correlation at E17.5, (r=−0.658, df=39, p=2.99*10^−6^) (**Fig. 8E**), but not at E18.5 (pooled r=−0.265, df=39, p=0.0946) (**Fig. 8F**), resulting at a significantly stronger correlation at E17.5 than at E18.5 (Z=−2.26, p=0.0244).

**Figure 8.**
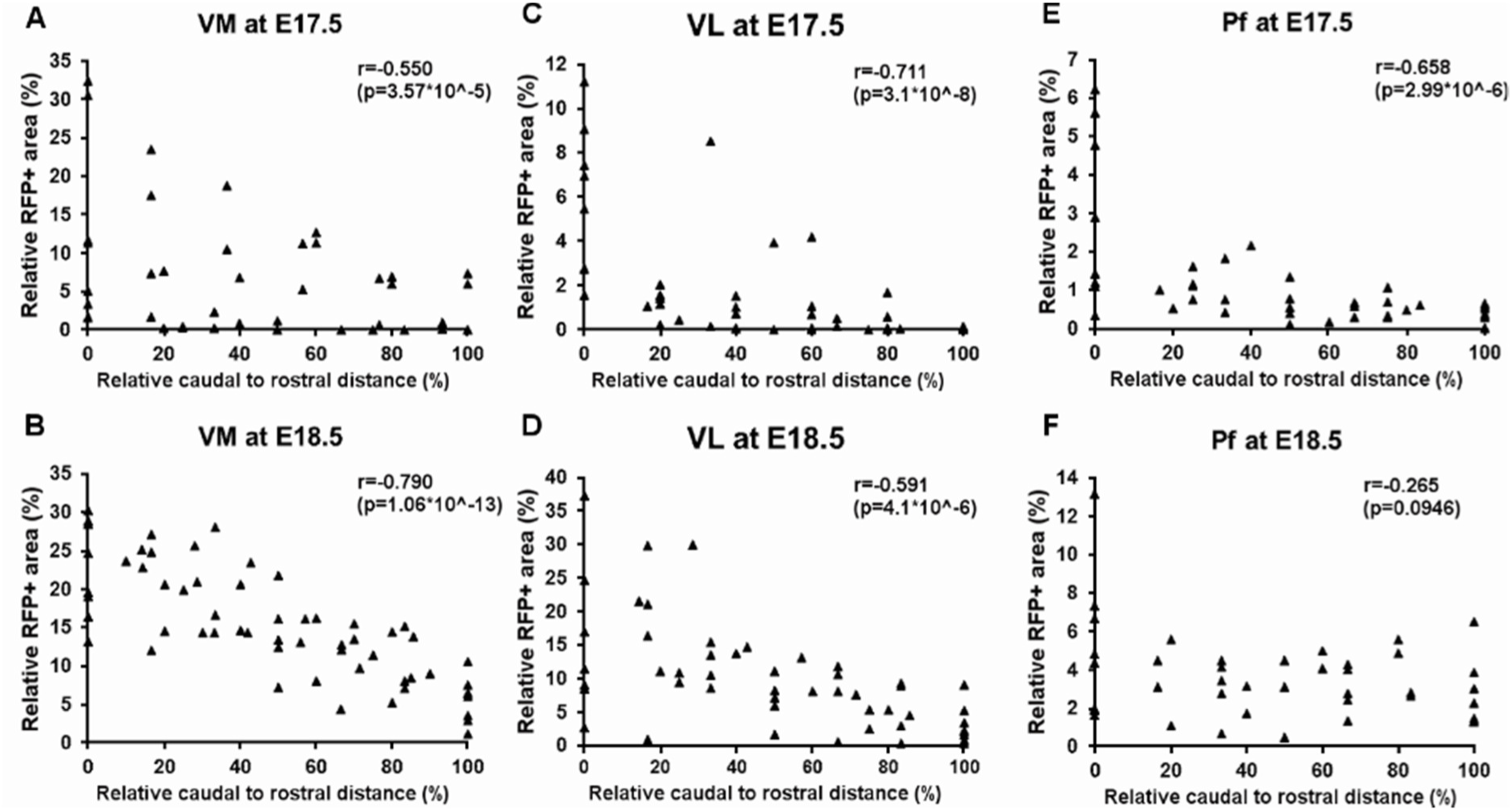
RFP signal quantification from caudal to rostral in the thalamic nuclei. The relative amount of RFP^+^ fibers (in %) is plotted against the relative rostral to caudal distance (in %) in VM, VL and Pf. A-B) VM at E17.5 (n=48) and E18.5 (n=57), respectively C-D) VL at E 17.5 (n=46) and E18.5 (n=52), respectively E-F) Pf at E17.5 (n=39) and E18.5 (n=39), respectively. r = Spearman’s rho. Pf = Parafascicular nucleus; VL = ventrolateral nucleus; VM = ventromedial nucleus

### Boutons and synapses

So far, we identified the location of RFP^+^ axons and the growth of the CbT in the dorsal thalamic complex. We next evaluated whether RFP^+^ axons formed synaptic contacts using confocal microscopy and FoxP2-stained tissue (**Fig. 6F, 9A**). Whereas our DAB and immunofluorescent staining of E17.5 tissue did not provide any hint for RFP^+^ bouton-like varicosities in thalamus (data not shown), at E18.5 RFP^+^ axons show morphological characteristics of pre-synaptic terminal formation throughout the thalamic complex (**Fig. 6F-H**; **Fig. 9B-I**). To confirm the subcortical origin of these terminals we co-labelled sections with vGluT2, i.e. a synaptic marker that separates subcortical axons from vGluT1-positive cortical layer VI axons (Fremeau et al., 2001; Hisano et al., 2002; Kaneko and Fujiyama, 2002). Stacks of deconvolved, high-magnification confocal images of VL (**Fig. 9B-E**), VM (**Fig. 9F-I**) and Pf (Fig. 9J-M) tissue confirmed the colocalization of vGluT2^+^ and RFP^+^ putative axon terminals. We found that the vGluT2-labelled RFP^+^ boutons in VL appear to have a larger volume (VL: 5.79±2.18 µm^3^, n=5 terminals; VM: 1.07±0.31 µm^3^, n=4 terminals) (**Fig. 9N**) although the difference was not significantly relevant (p=0.11, Mann Whitney test).

**Figure 9.**
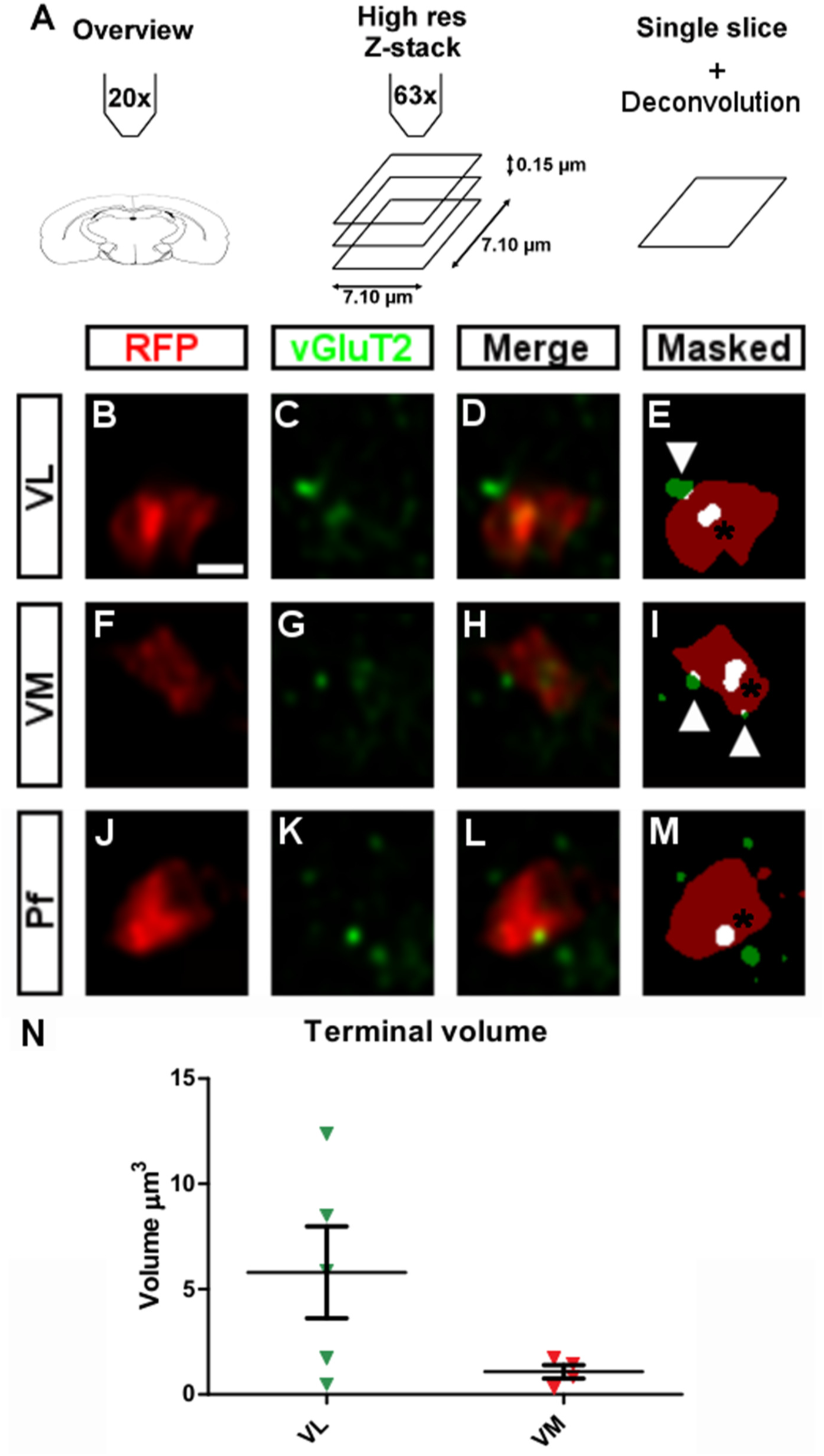
Putative boutons and colocalization with vGluT2. A) Schematic overview of the workflow. After taking an overview image at 20X, we acquired z-stacks at 63X and deconvolved the maximum projection, which is represented as a single optical slice. A user-defined threshold determined the overlap between fluorophores. RFP^+^ boutons were considered to colocalize with vGluT2 if the vGluT2-fluorescence overlapped completely in all dimensions. B-E) Example of a deconvolved image of a putative bouton in VL, with RFP in red (B), vGluT2 in green (C), colocalization of RFP and vGluT2 (D), and the result after thresholding (E). Asterisks indicate complete overlap, arrowheads indicate partial overlap, the latter of which is not considered for further analysis. F-I) as B-E for VM. J-M) as B-E for Pf. N) Terminal volumes (in µm^3^) in VL and VM. Scale bars: B-M = 1 µ m. Pf = Parafascicular nucleus; RFP = red fluorescent protein; vGluT2 = vesicular glutamate transporter 2; VL = ventrolateral nucleus; VM = ventromedial nucleus.

## Discussion

We describe the outgrowth of CN axons into the thalamic complex. Our data reveal that at E15.5 and E16.5 the CbT reached the prethalamus and from E17.5 onwards is detected throughout the developing thalamus. During these last days of the embryonic stage the thalamic nuclei that receive the densest CN axon projections in the adult stage, like VM, VL and Pf, were increasingly invaded by RFP^+^ axons. At E18.5 we found RFP^+^ axon varicosities that colocalized with vGluT2.

### Technical considerations

Using *Ntsr1-Cre/Ai14* brains for our current study allowed us to investigate the development of CbT axonal projections, but also resulted in RFP^+^ expression in other cell populations, some of which are known to innervate the thalamic complex. Apart from the CN labelling (see also(Houck and Person, 2015)), RFP^+^ neurons and axons were also readily identified in the deeper layers of the cerebral cortex, where *Ntsr1* is known to be expressed by a subpopulation of L6 pyramidal cells (Gong et al., 2007; Olsen et al., 2012). As we have shown in our analysis the corticothalamic (CT) tract, which contains the L6 fibers, and the CbT tract are positioned differently in the embryonic mouse brain (**Fig. 3**). Also the time of thalamic invasion by the CT and CbT fibers is different: our data reveal that already from E17.5 RFP^+^ fibers start invading the thalamic complex, which precedes the innervation of sensory and motor thalamic nuclei by CT fibers, which has been shown to occur earliest from E18.5 (as reviewed by (Grant et al., 2012)). Finally, the nuclei that are innervated by CT and CbT tracts differ; i.e. CT fibers initially innervate the ventrobasal nuclei and appear to diverge from their onwards (Jacobs et al., 2007), whereas the CbT arrives in the ventromedial nucleus.

We also found sparse labelling of somata in several other brain regions, which has implications for the interpretability of our data on thalamic innervation in the embryonic stages. Of these regions we found that mRF cells that project to the thalamus are located more dorsally than where we observed Ntsr1^+^ cells (Newman and Ginsberg, 2008). Also the hypothalamic Ntsr1^+^ cells, which in principle could be part of the subset of thalamus-projecting neurons, we conclude that the proportion of RFP^+^ fibers in the VL, VM and Pf nuclei is limited (Vertes et al., 1995; Shimogawa et al., 2015). In addition, we showed in E18.5 tissue that labelling CbT axons via IUE transfection of CN neurons yielded a similarly labeled CbT tract with a similar projection pattern as our *Ntsr1-Cre/Ai14* embryos, further corroborating our assumption that the bulk of RFP-signal in the embryonic thalamus originates from the CbT tract.

### Development of the CbT in mouse embryos

In the present study, we investigated the anatomical development of the CbT in mouse embryos aged E15.5 to E18.5 (Fig. 10) and found that prior to the entry of the thalamus, the CbT tract appears to stall its growth and reside in the prethalamus between E15.5 and E16.5. Several other thalamic afferents reveal such a ‘pause’ or waiting period, such as the trigeminal projection to the sensory VPM (Kivrak and Erzurumlu, 2013) and CT pathways (Jacobs et al., 2007). For the CbT this waiting period could come about due to several reasons: *i*) to gain axonal energy supplies to allow axonal branching which could demand extra mitochondria to be produced (Sheng, 2017); *ii*) although most of not all synaptic targets of CN axons are already born, CN invasion could be halted until the neuronal migration nears completion (Altman and Bayer, 1989) and *iii*) CbT fibers, which originate from CN neurons born sequentially (Leto et al., 2015), regroup in the prethalamus before entering the thalamic complex – a phenomenon described for the CT tract (Grant et al., 2012). Also the mechanisms that facilitate the CbT waiting period remains to be elucidated. We hypothesize that the CbT tract is stalled temporarily by chemical signaling. Possible candidate signaling cascades are semaphorin-based, (Kolk et al., 2009; Deck et al., 2013), albeit that numerous other molecules and receptors can potentially also influence the development of the CbT (Bibollet-Bahena et al., 2017).

**Figure 10.**
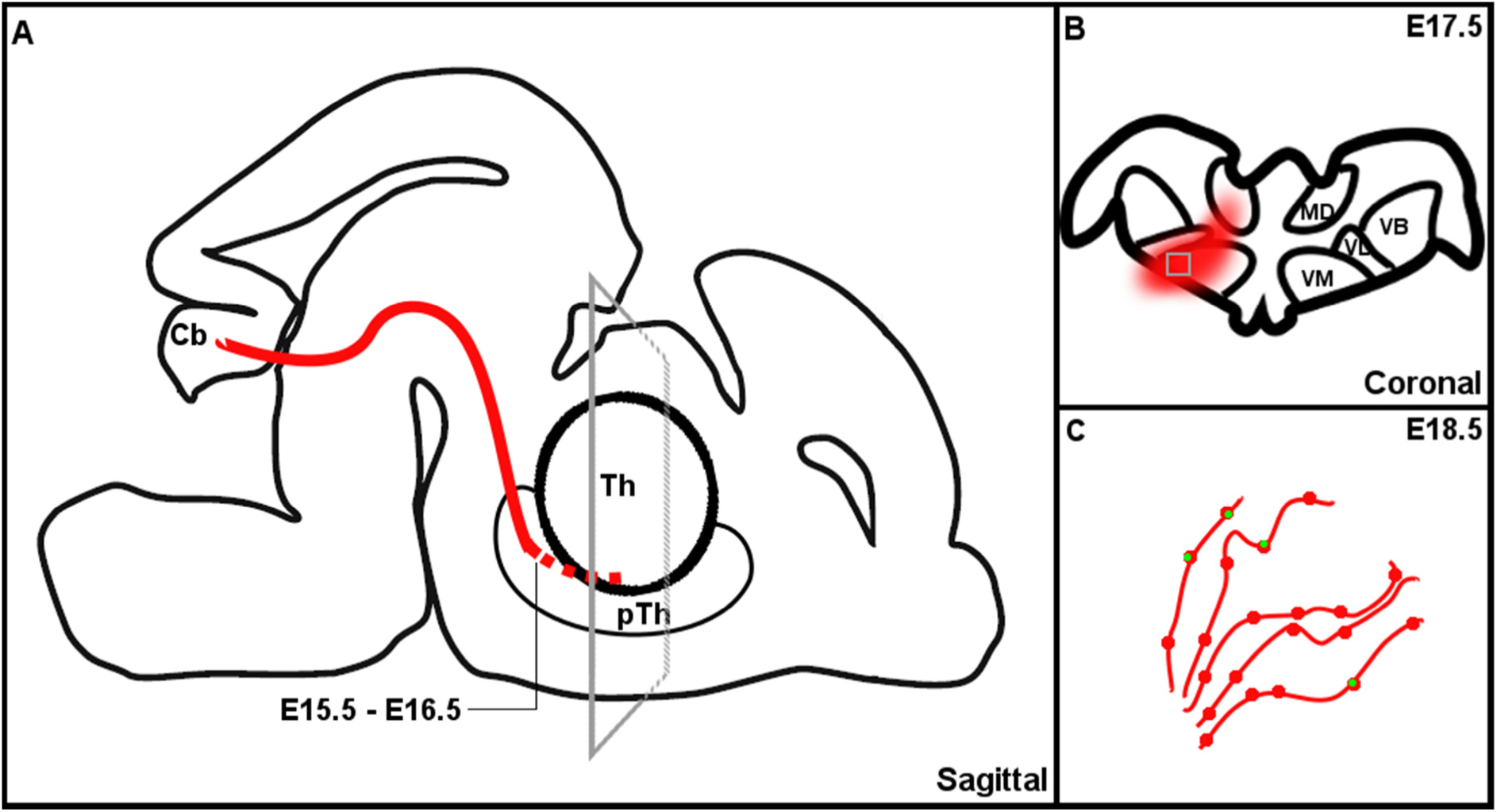
Schematic representation of the development of the cerebellothalamic tract. A) Sagittal representation of the development of the cerebellothalamic tract. B) Coronal representation of inset in A, showing schematic representation of the invasion of cerebellothalamic fibers into the thalamus. C) Zoom in of inset in B, showing a schematic representation of the appearance of vGluT2 positive terminals at E18.5.Cb = cerebellum; pTh = prethalamus; Th = thalamus; vGluT2 = vesicular glutamate transporter 2.

Upon entry in thalamus, CbT fibers appeared to swiftly locate the VM and VL nuclei, which are also at adult stages their prime target nuclei (Teune et al., 2000; Gornati et al., 2018). We found that the most caudal portions of VM and VL nuclei showed more RFP^+^ labeling than the rostral portions, indicating that the afferents arise from caudal, which matches with the position of the CbT tract. The fact that the rostro-caudal gradient disappeared in Pf could suggest that CbT innervations to Pf has stabilized already at E18.5 or that also fibers that arrive in a different orientation start to innervate this nucleus, like the hypothalamic-thalamic fibers (Shimogawa et al., 2015). Further investigations of the various thalamic afferents shall reveal more insights in how their growth is organized.

### Timing of invasion in the thalamus

The thalamic invasion by CbT fibers starts from approximately E17.5. This is relatively late compared to other subcortical thalamic afferents. Both primary sensory afferents and serotonergic afferents start invading the thalamus prior to CbT fibers. The retinogeniculate pathway is present in the LG as early as E15.5 in mouse (Moreno-Juan et al., 2017) and the trigeminothalamic pathway invades the thalamus between E14 and E17 in mouse(Jones, 2007; Kivrak and Erzurumlu, 2013). Serotonergic afferents from the brainstem enter the thalamus at approximately E16 in rat (Lauder et al., 1982). As has been proposed before based on studies in the opossum, these ascending brainstem fibers may provide a passage for the CbT fibers through the mesencephalon (Martin et al., 1987), but whether there is active interaction between these thalamic afferents remains to be elucidated

It remains speculative what role the formation of CbT connectivity during late embryonic stages has, not in the least because the cerebellar cortex is developing relatively late and its impact on the CN neurons is rudimentary at best in the antenatal period (Gardette et al., 1985). One option is that CbT fibers are present in the thalamic complex to interact with thalamic neurons and guide their axonal growth. The invasion of thalamocortical (TC) fibers into the cortical plate starts from E18.5-P0.5, but already before that the activity levels of thalamic neurons are thought to direct the axonal growth speed (Uesaka et al., 2007; Matsumoto et al., 2016; Moreno-Juan et al., 2017). Thus, since the CbT tract seems to have established connections with the thalamus starting at E18.5, there is ample time for the cerebellar output to affect thalamo-cortical development.

Both the thalamocortical system and the cerebellum have been implicated to undergo critical periods (Kivrak and Erzurumlu, 2013; Wang et al., 2014). Disruptions of the cerebellar development are suggested to cause disrupted behavioral traits in the motor and non-motor domain, linking to various neurodevelopmental disorders and psychiatric diseases, like autism spectrum disorder and schizophrenia (Wang et al., 2014; Sathyanesan et al., 2019). These neurological conditions are thought to be related to malformations in the neuronal wiring caused by adverse early life events (Di Martino et al., 2014; Nair et al., 2015) and our current study suggests that in the murine brain cerebellar aberrations could possibly derail thalamocortical wiring from E17.5 onwards, which translates to early fetal stages in human development. Future experiments can investigate the impact of cerebellar abnormalities on TC development and cortical maturation.

## List of abbreviations

CbT: cerebellothalamic tract
CbX: decussation of the cerebellothalamic tract
CN: cerebellar nuclei
CP: Caudate Putamen
CT: CT
Ctx: cerebral cortex
Th: thalamus
fr: fasciculus retroflexus
Hb: Habenula
Hc: hippocampus
Hyp: Hypothalamus
IC: inferior colliculus
Int: interposed nucleus of the cerebellum
LCN: lateral nucleus of the cerebellum
LG: lateral geniculate nucleus of the thalamus
dLGN: dorsal lateral geniculate nucleus
LP: lateral posterior nucleus of the thalamus
MCN: medial nucleus of the cerebellum
MD: mediodorsal nucleus of the thalamus
MG: medial geniculate nucleus of the thalamus
ML: midline nuclei
mRF: medullary reticular formation
mt: mamillothalamic tract
NHS: normal horse serum
PBS: phosphate buffer saline
PBSGT: phosphate buffer saline with gelatin and triton
Pf: parafascicular nucleus of the thalamus
POm: Posterior complex of the thalamus
pSAO: percentage of summed area occupied by above threshold RFP signal
Pt: pretectum
RFP: red fluorescent protein
RN: red nucleus
rpm: rounds per minute
Rt: reticular nucleus of the thalamus
SC: superior colliculus
TC: thalamocortical
THF: tetrahydrofuran
VB: ventrobasal complex of the thalamus
VL: ventrolateral nucleus of the thalamus
VM: ventromedial nucleus of the thalamus
pTh: prethalamus
ZI: zona incerta

## Acknowledgements

The authors thank Elize Haasdijk, Erika Sabel-Goedknegt, Mandy Rutteman, Martin van Royen, Gert-Jan Kremers and Alex Nigg for technical assistance, Rosa Lahikainen, Sverrir Sigurdsson, Bas van Hoogstraten, Carmen B. Schäfer, Oscar Eelkman Rooda and all other members of the Hoebeek team for constructive discussions. This work was supported by NWO-ALW VIDI 016.121.346, Zon-MW TOP GO 91210067 and a generous contribution of the C.J. Vaillant fund to F.E.H. NWO-ALW VICI and a Utrecht University facility grant to R.J.P.

